# Ecological drivers of sustained enzootic yellow fever virus transmission in Brazil, 2017-2021

**DOI:** 10.1101/2022.10.19.512702

**Authors:** Natalia Ingrid Oliveira Silva, Gregory F Albery, Matheus Soares Arruda, Gabriela Garcia Oliveira, Thaís Alkifeles Costa, Érica Munhoz de Mello, Gabriel Dias Moreira, Erik Vinicius Reis, Simone Agostinho da Silva, Marlise Costa Silva, Munique Guimarães de Almeida, Daniel J. Becker, Colin J. Carlson, Nikos Vasilakis, Kathryn Hanley, Betânia Paiva Drumond

## Abstract

Beginning December 2016, sylvatic yellow fever (YF) outbreaks spread into southeastern Brazil, and Minas Gerais state experienced two sylvatic YF waves (2017 and 2018). Following these massive YF waves, we screened 187 free-living non-human primate (NHPs) carcasses collected throughout the state between January 2019 and June 2021 for YF virus (YFV) using qPCR. One sample belonging to a *Callithrix*, collected in June 2020, was positive for YFV. The viral strain belonged to the same lineage associated with 2017-2018 outbreaks, showing the continued enzootic circulation of YFV in the state. Next, using data from 781 NHPs carcasses collected in 2017-18, we used generalized additive mixed models (GAMMs) to identify the spatiotemporal and host-level drivers of YFV infection and intensity (an estimation of genomic viral load in the liver of infected NHP). Our GAMMs explained 65% and 68% of variation in virus infection and intensity, respectively, and uncovered strong temporal and spatial patterns for YFV infection and intensity. NHP infection was higher in the eastern part of Minas Gerais state, where 2017-2018 outbreaks affecting humans and NHPs were concentrated. The odds of YFV infection were significantly lower in NHPs from urban areas than from urban-rural or rural areas, while infection intensity was significantly lower in NHPs from urban areas or the urban-rural interface relative to rural areas. Both YFV infection and intensity were higher during the warm/rainy season compared to the cold/dry season. The higher YFV intensity in NHPs in warm/rainy periods could be a result of higher exposure to vectors and/or higher virus titers in vectors during this time resulting in the delivery of a higher virus dose and higher viral replication levels within NHPs. Further studies are needed to better test this hypothesis and further compare the dynamics of YFV enzootic cycles between different seasons.

**Author Summary:** In 2017 and 2018 massive sylvatic yellow fever (YF) outbreaks took place in Minas Gerais Brazil. To investigate yellow fever virus (YFV) circulation following these massive outbreaks, we investigated samples from 187 free-living non-human primate (NHPs) collected between January 2019 and June 2021. One sample belonging to a *Callithrix*, collected in June 2020 was positive for YFV. This virus was closely related to YFV from previous outbreaks (2017-2018) showing the continued enzootic circulation of YFV in the state. Next, we investigated the drivers of YFV infection and intensity (an estimation of viral load in each infected NHP) during the 2017-18 outbreaks. The odds of YFV infection in NHPs were lower in urban areas compared to rural ones as expected in sylvatic cycles. There were strong temporal and spatial patterns were observed for YFV infection and intensity, especially in the eastern part of Minas Gerais state. The higher YFV infection and intensity observed during the warm/rainy season (as opposed to the cold/dry one) could be related to higher exposure to vectors and/or higher virus titers in vectors during this time. The possible delivery of a higher virus dose in NHPs could lead to higher viral replication levels within NHPs explaining the higher intensity of infection during warm/rainy season. Further studies are needed to better test this hypothesis and further compare the dynamics of YFV enzootic cycles between different seasons.

## Introduction

Mosquito-borne yellow fever virus (YFV; family *Flaviviridae*, genus *Flavivirus*) originated in tropical and subtropical areas of Africa and was introduced to South America after European colonization (1). YFV was almost certainly transported to the Americas via infected humans or *Aedes aegypti* mosquitoes, and while the urban cycle of yellow fever (YF) in South America continued to be maintained by *Ae. aegypti* and *Ae. albopictus* (2–4), the virus also spilled back into non-human primates (NHPs) and arboreal, hematophagous, primatophilic mosquitoes of the genera *Haemagogus* and *Sabethes* (3). In Brazil, the urban cycle of YF has not been documented in Brazil since 1942 (3,4). However, sylvatic YF cycles have persisted within the Brazilian Amazon Basin (2–4) with periodic expansions towards the Central and Southern regions of the country during the warm and rainy seasons (December to May in Southeast Brazil) (3–5). The YF surveillance program in Brazil focuses on the investigation of suspected human cases, NHP epizootics, and entomological surveillance (6), with annual reports spanning the typical transmission seasons of the disease (July to June of the following year).

In recent decades, many experts have warned that new urban outbreaks could arise from sylvatic YFV cycles in South America and deplete the limited supply of YFV vaccine (1,7–10). Although these fears have not yet been realized in their entirety, between 2016 and 2018 Brazil faced the largest sylvatic YF outbreak since the eradication of urban YF. The virus spread rapidly in the Southeast region of the country, reaching densely populated areas, causing thousands of cases and deaths in both NHPs and humans, and raising the specter of full-blown urban transmission mediated by *Ae. aegypti* (3,4,11–15). Between 2016 - 2019, 2,251 YF human cases were confirmed in the Southeast region of Brazil, with 772 deaths reported (3,4,16). Additionally, during this period, approximately 15,000 NHPs deaths were reported in the country, with the confirmation of at least 1,500 cases linked to YFV (3). YF cases waned in Minas Gerais state during 2018; however, the virus has been spreading southwards since 2019, affecting humans and NHPs in São Paulo, Paraná, Rio Grande do Sul, and Santa Catarina states (3,16–18). Since then, YF epizootics have been confirmed outside the Brazilian Amazon Basin during the YF monitoring periods of Jul 2020/Jun 2021 (Midwest: Goiás state and Federal District; Southeast: São Paulo state, and South region: Paraná, Santa Catarina, and Rio Grande do Sul states) (19), and Jul 2021/Jun 2022 (South: Santa Catarina, and Rio Grande do Sul states, and Southeast region: Minas Gerais state) (20).

YF epizootics in NHPs often precede and overlap the occurrence of human cases (3,21), as was observed in the recent outbreaks in Brazil (22,23). Although neotropical NHPs are highly susceptible to YFV infection and sentinels for viral circulation, the susceptibility of different NHP genera and species to YFV infection is still poorly characterized (3). Some insights have been provided by investigations of NHPs during the recent epizootics in Brazil. *Callithrix* spp. (marmoset) specimens showed lower genomic viral loads (estimated by the threshold for cycle quantification value (Cq) in RT-PCR assays) compared to *Alouatta* spp. (howler monkey), *Callicebus* spp. (titi monkey) (11,24–26), and *Sapajus* spp. (capuchin monkey) specimens (24–26). Histopathological and immunohistochemical analyses also suggested differences among NHP genera in susceptibility to YFV infection (24,26) and mortality rates to the virus (26). *Alouatta* was the most affected group of NHP, with the highest proportional YF-related mortality rate. *Callicebus, Alouatta*, and *Sapajus* had similar genomic YFV loads, but the highest proportional mortality rate was observed in *Alouatta* infections. *Callithrix* had the lowest genomic viral loads and proportionally lowest mortality rates (26).

NHP population dynamics and susceptibility to viral infection play key roles in viral transmission and spillover (27), as do the ecological dynamics and interactions between YFV and its vectors, hosts, and abiotic factors (4,28). Not surprisingly, previous studies have demonstrated that NHP immunity to YFV and infection prevalence are key missing variables in predictive models, representing a significant gap in our understanding of YFV transmission that impairs our ability to forecast spillover (28). The recent reemergence of sylvatic YF in Brazil into highly densely populated regions with low vaccination coverage in Southern Brazil clearly exposes the inadequate monitoring of YFV circulation in mosquitoes and NHPs and the underestimation of the risk of YF beyond the endemic areas. Although the seasonality of YF is well defined, little is known about the environmental, climatic, vector, and host drivers involved (29). Some studies have already shown that some abiotic variables such as vegetation, rainfall, temperature, and human population size are important drivers in the transmission of YFV (29). Coupled field surveillance and modeling approaches could provide crucial advances in containing, and possibly averting, future YFV outbreaks (2,4).

Here, we investigated the occurrence of YFV in NHPs from Minas Gerais state between 2019-2021, following the outbreaks in 2017 and 2018. We used generalized additive mixed models (GAMMs) to identify the spatiotemporal drivers of YF infection and intensity in NHPs, specifically testing the hypotheses that: (i) NHP and human YFV infection prevalence would be synchronized, with peaks in NHP infections preceding those in humans, as transmission from NHP to humans is thought to occur but the reverse is not; and (ii) that YFV infections of NHP would show spatial structure across the state of Minas Gerais and among different land cover types.

## Materials and Methods

### Study area and samples

The samples and data analyzed in this study were collected from Minas Gerais, the epicenter of the YF outbreak in 2017 and 2018 in Brazil, with records of 1,002 cases and 340 deaths of humans, between July 2016 – June 2018 (30). Minas Gerais borders the states of São Paulo, Rio de Janeiro, and Espírito Santo in Southeast Brazil (Fig 1a). It has a total population of 21,168,791 inhabitants, with approximately 85% living in urban areas and the remaining population living in rural areas (31). The state is mainly covered by three biomes: Atlantic Forest, Cerrado (Brazilian Savana), and Caatinga, where different genera of NHPs can be found, including *Alouatta, Callithrix, Sapajus, Callicebus*, and *Brachyteles* (woolly spider monkeys) (32–34).

**Figure 1:**
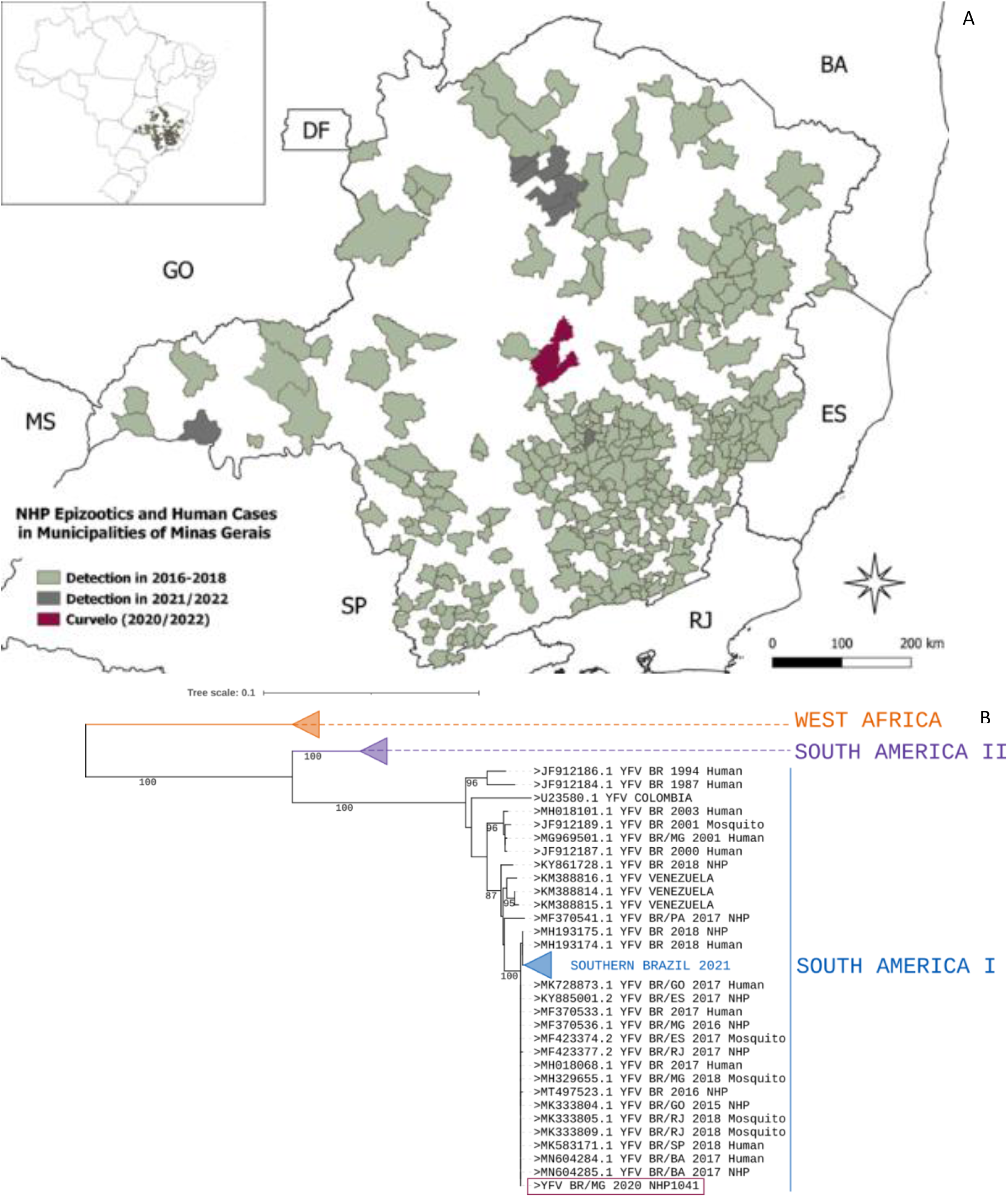
Circulation of yellow fever virus in Minas Gerais, 2016 to 2021. A: Map of Minas Gerais showing the location of Curvelo, where yellow fever virus was detected in a *Callithrix* sp specimen, in 2020 (purple), municipalities that had cases of yellow fever affecting human and or non-human primate, 2016-2018 (green), and municipalities that had yellow fever epizootics confirmed from Jul 2020 to Jun 2022 (grey). The inset map displays the location of Minas Gerais in Southern Brazil. B: Phylogenetic tree reconstructed with using partial envelope sequence of yellow fever virus (1000nt). The sequence of yellow fever virus detected in a *Callithrix* sp specimen (NHP1041), in 2020, from Curvelo is shown in purple. Some branches were collapsed for clarity. The phylogenetic tree was reconstructed using the maximum likelihood method, nucleotide substitution model TN93 +I, with 1000 bootstrap replicates, using PhyML. The final tree was edited and visualized using iTOL.

From January 2019 to July 2021, a total of 543 carcasses of free-living NHPs were collected in Minas Gerais by health surveillance agents as part of the YF surveillance program (mean of 30 carcasses/month). The number of NHP carcasses sampled was 3.8 times lower compared to December 2016 up to December 2018 (mean of 86 carcasses/month), when YF outbreaks were reported in the region. The NHP carcasses were frozen (−20 °C) and forwarded to the Laboratory of Zoonosis of Belo Horizonte/Minas Gerais (LZOON-MG). At LZOON-MG, it was only possible to harvest biological specimens (liver and lungs) from 187 individuals (2019: n=123; 2020: n=41; 2021: n=23), due to the carcasses’ preservation status. In 2019, only liver samples were harvested from carcasses. Starting in 2020 lung samples were also harvested from some carcasses, with the parallel purpose of investigating infection by SARS-CoV-2 (35); lung samples were also tested for YFV whenever available. Liver and lung samples were preserved in RNAlater solution (Ambion/USA), sent to the Virus Laboratory at Universidade Federal de Minas Gerais, and held at −20 °C until further use. The study was approved by the Institutional Animal Care and Use Committee of Universidade Federal de Minas Gerais under protocol CEUA: 98/2017 (approval date: June 26^th,^ 2017).

NHP carcasses were identified to the genus level by the veterinary team at LZOON-MG using morphological criteria. Each carcass was georeferenced by the location of sampling and geographically classified by land use (urban, rural, urban-rural interface). Land use was classified using satellite imagery (36) as i) urban areas (built-up areas, cities, towns, villages, or isolated areas with intense human occupation and urban development); ii) urban-rural interface (transition between urban and rural areas delimited by 2.0 kilometers on the outskirts of urban areas towards the rural/sylvatic environment); and iii) rural area (sylvatic areas that covered areas located outside the urban and urban-rural transition areas) (11,37).

### Molecular screening for YFV in NHP samples

Total RNA was extracted in batches of a maximum of 15 samples at a time (liver for all NHPs and lung samples as described above), plus one negative extraction control (nuclease-free water), using the RNeasy Minikit (Qiagen, USA). To determine the suitability of the RNA, samples were submitted to a one-step real-time polymerase chain reaction (RT-qPCR), using the GoTaq 1-Step RT-qPCR System (Promega, USA) and primers targeting the gene coding for the endogenous control β-actin. All RNA samples were positive for the endogenous control (11,38). RNA samples and the negative extraction controls were screened in duplicate to investigate the presence of YFV RNA by RT-qPCR (GoTaq Probe 1-Step RT-qPCR System-Promega, USA), with primers and probes targeting the 5’-noncoding regions of the YFV genome (39). Non-template, negative extraction, and positive controls (RNA from the YFV 17DD vaccine, provided by Bio-Manguinhos/FIOCRUZ) were used in every assay.

Samples were considered positive for YFV when they presented a threshold for cycle amplification (Cq) ≤ 37 in duplicate (11). Samples with Cqs >37 and ≤40 were considered indeterminate and retested, and samples with Cqs > 40 were considered negative. When YFV RNA was detected, the sample was retested without the presence of reverse transcriptase to exclude amplicon carryover. In the case of a YFV positive sample, the clarified supernatant from the macerated sample was used to infect C6/36 and Vero cells, followed by consecutive blind passages attempting viral isolation (38). The supernatant of infected cells was used for RNA extraction (Qiamp Viral RNA Mini Kit, Qiagen) and tested by RTqPCR as described above. In addition, RNA from the positive sample was used as a template to generate amplicons covering part of the envelope gene of YFV using different primer combinations (YFV_500_3_left/right, YFV_500_4_left/right, YFV_500_5_left/right, and YFV_500_6_left/right) as previously described (22) using the GoTaq Probe Master Mix (Promega, USA). The amplicons were purified and sequenced with the ABI3130 platform (Applied Biosystems, USA). The consensus sequences were generated (<u>Chromas software version 2.6.6</u>), aligned with MAFFT (mafft.cbrc.jp/alignment/server/) and used for phylogenetic reconstruction, using the maximum likelihood method, using the nucleotide substitution model that best fitted the data (TN93 +I), with 1000 bootstrap replicates, using PhyML (40). The final tree was edited and visualized using iTOL (41).

### Generalized additive mixed models

Generalized additive mixed models (GAMMs) were used to identify the drivers of YFV infection and intensity because these models allow for non-linear trends with time, space, and abiotic drivers. All models were fitted in R version 4.1.1 (42), using the ‘mgcv’ package (43). Response variables included binary qualitative liver RTqPCR results, denoting the presence or absence of infection (“infection”), and RTqPCR Cq threshold values (inversely proportional to genomic YFV load in the liver of NHPs) that represent the intensity of YFV in infected individuals (“intensity”).

To conduct the analysis, we initially combined our data from 2019-2020 with previously published data from molecular YFV screening in 781 NHP carcasses from 2017 to 2018 in Minas Gerais (11,38). However, given the extreme low positive rate for YFV in 2019-2021 (see Results), our models only used data from 2017 and 2018. Intensity models (i.e., RTqPCR Cq values) only included samples that were positive for YFV. RTqPCR Cqs threshold values were inverted so that larger values denoted greater intensity, with values then scaled to have a mean of 0 and a standard deviation of 1. Infection and intensity models used a binomial and Gaussian specification for the response, respectively.

GAMMs were used to investigate several key hypotheses about the drivers of YFV infection and intensity in NHPs outlined above. First, the epizootics were divided into two periods pre-and post-day 220 (corresponding to Aug 8^th^ 2017, which was the nadir of prevalence through the cold/dry season), and an interaction between the spatial effect and the epizootics (first or second) was fitted to examine whether they differed in their geographic distributions. Finally, temporal patterns and changes in infection and intensity in our sample of NHPs were compared with total confirmed YF cases in human and NHPs in Minas Gerais state, using data from Information System for Notifiable Diseases (SINAN) the Brazilian Ministry of Health’s official yellow fever surveillance program. Data regarding confirmed YF cases in human and NHPs were received from SINAN (July 8^th^ 2021) and date of epizootic occurrence for NHPs, or date of symptom onset for humans were used.

Explanatory variables in our models included: a smoothed effect of Julian date with an adaptive smooth to examine temporal patterns; a latitude-by-longitude smoothing term to examine spatial patterns; a three-level factor to denote sample land use (urban, urban-rural interface, and rural); a three-level factor to denote carcass preservation quality (good, intermediate, and bad); and a three-level factor to denote the genus of NHP sampled (*Alouatta, Callicebus*, and *Callithrix*). We also attempted to fit the effects of local population density and gross domestic product (data from Brazilian Institute for Geography and Statistics – Instituto Brasileiro de Geografia e Estatística/IBGE), but neither improved the model fit or showed significant effects, and both required a substantial reduction in sample size due to missing data, so we opted not to include them in our final models. For each effect in the GAMMs, contribution to model fit was calculated by examining their effect on the Spearman rank correlation between the models’ observed and predicted values (which we call R). This calculation involved removing a variable from the model’s predictions and calculating the correlation of the new predictions with the observed data. Removing more important variables has a greater effect on the model’s fit than other, less important ones. These are then expressed as a proportion of the model’s overall fit to give a representation of each variable’s importance in the model. All code associated with this analysis is available at https://github.com/viralemergence/minas-gerais-yfv.

## Results and Discussion

### YFV enzootic circulation in in NHP from Minas Gerais between 2019-2021

From January 2019 to July 2021, we screened 187 free-living NHP carcasses collected during the warm/rainy (December to May) and cool/dry periods (June to November) of 2019 - 2021. Both lungs and liver were screened for YFV where available; liver was available for all carcasses, and for 44 carcasses lung specimens were also available. Carcasses were collected from 70 municipalities throughout the state but mostly originated from three regions, the Metropolitan (n=101, 54.0%), Zona da Mata (n= 21, 11.2%), and Triângulo/Alto Paranaíba (n=20, 10,7%). The great majority of these carcasses (93.6%) were *Callithrix* from urban areas (Table 1).

**Table 1.** Non-human primate carcasses collected from 2019 to 2021 in Minas Gerais and screened for yellow fever virus

| Non-Human Primate |  | year |  |  | Total |
| --- | --- | --- | --- | --- | --- |
|  |  | 2019 | 2020 | 2021 |  |
| Period* | warm/rainy | 51 | 20 | 12 | 82 (43.8%) |
|  | cool/dry | 73 | 21 | 11 | 105 (56.2%) |
| Area | Urban | 95 | 31 | 23 | 148 (78.7%) |
|  | Urban-rural interface | 5 | 2 | 0 | 7 (4.2%) |
|  | Rural | 24 | 8 | 0 | 32 (17.1%) |
| Taxon | *Callithrix* spp. | 114 | 39 | 22 | 175 (93.6%) |
|  | *Alouatta* spp. | 0 | 1 | 1 | 2 (1.1%) |
|  | *Cebidae* spp. | 9 | 1 | 0 | 10 (5.3%) |
| Samples** | Liver | 124 | 41 | 23 | 187 |
|  | Lung | 0 | 21 | 23 | 44 |

\* warm/rainy period: from December to May, cool/dry period: June to November.
Samples\*\*: Liver samples were available and analyzed for all non-human primates' carcasses. For all carcasses (n=187) liver samples were analyzed, but in some cases (n=44), besides liver, lung was available and also analyzed.

Between 2019 and 2020, one out of 44 lung samples tested positive for YFV. This positive sample was from a *Callithrix* specimen collected during the cold/dry season in June 2020, in the central area of Minas Gerais (Fig 1a). None of 187 liver samples tested positive during this period. Three independent RNA extraction and RT-PCR experiments were performed using lung and liver samples from the YFV positive carcass with similar results. DNA template carryover during experimental procedures was ruled out by running PCR without previous reverse transcription. Attempts at viral isolation from the positive lung sample were not successful after five blind passages in C6/36 or Vero cells, since no YFV RNA was detected in supernatant from infected cells. The high Cq (= 38.8) observed in RTqPCR of the positive sample, indicated a very low genomic load reflected in a low viral load impairing viral isolation. A nucleotide sequence of partial envelope gene (1113 nt, Genbank accession number OP589291) of YFV confirmed the detection of wild-type YFV RNA. Phylogenetic analyses demonstrated that this YFV strain clustered within a monophyletic clade of South America genotype I, together with other YFV isolates from the recent outbreaks (2016 to 2019) in Brazil (Fig 1b).

The YFV-positive *Callithrix* specimen was collected in the rural area of the municipality of Curvelo, central Minas Gerais state, an area that was heavily affected by YF in 2017 and 2018 and from which NHP epizootics have been reported since (44,45). The molecular and epidemiological data point to the maintenance of YFV lineage related to 2017 and 2018 outbreaks in the enzootic cycle in Minas Gerais state up to 2020. This lineage has been proven to have undergone cryptic circulation prior to the outbreak by the end of 2016 (22,38,46), and to have persisted in Southern Brazil at least until 2021 (6,38,46-48).

Despite the significant decrease in the magnitude of NHP epizootics from the first two YF waves (2016/2017 and 2017/2018) to subsequent warm/rainy seasons, our data and data from official surveillance systems reinforce the conclusion that YFV circulates continuously in an enzootic circulation in Minas Gerais. During the last YF surveillance period (July 2021-June 2022), eight municipalities in the state, located in areas covered by Cerrado biome (Brazilian Savana) confirmed epizootics caused by YFV (44,45,49). The very low prevalence of YFV in NHPs from 2019 to 2021 observed here may reflect a real reduction in transmission or may be driven by the fact that the majority of NHP carcasses were collected in urban areas (78.73%). Below we demonstrate that odds of YFV infection in NHPs from urban areas are significantly lower than rural-urban interface or rural areas. These carcasses represent a convenience sample obtained through the YF surveillance system. The shift in the distribution of submitted NHP carcasses from a more even spread across land cover types to an almost exclusive representation of urban areas may be driven by the public concern about the recent circulation of YFV in densely populated areas in southeastern states of Brazil (11-15,30).

### Drivers of YFV infection and intensity during epizootics

For YFV data collected from NHPs in Minas Gerais from 2017-2018, our GAMMs explained 65% and 68% of the variation in virus infection and intensity, respectively. Model coefficients and associated p values for infection and intensity models are listed in Table 2 with pseudo-R^2^ values in Table 3 to illustrate model fit.

**Table 2:**
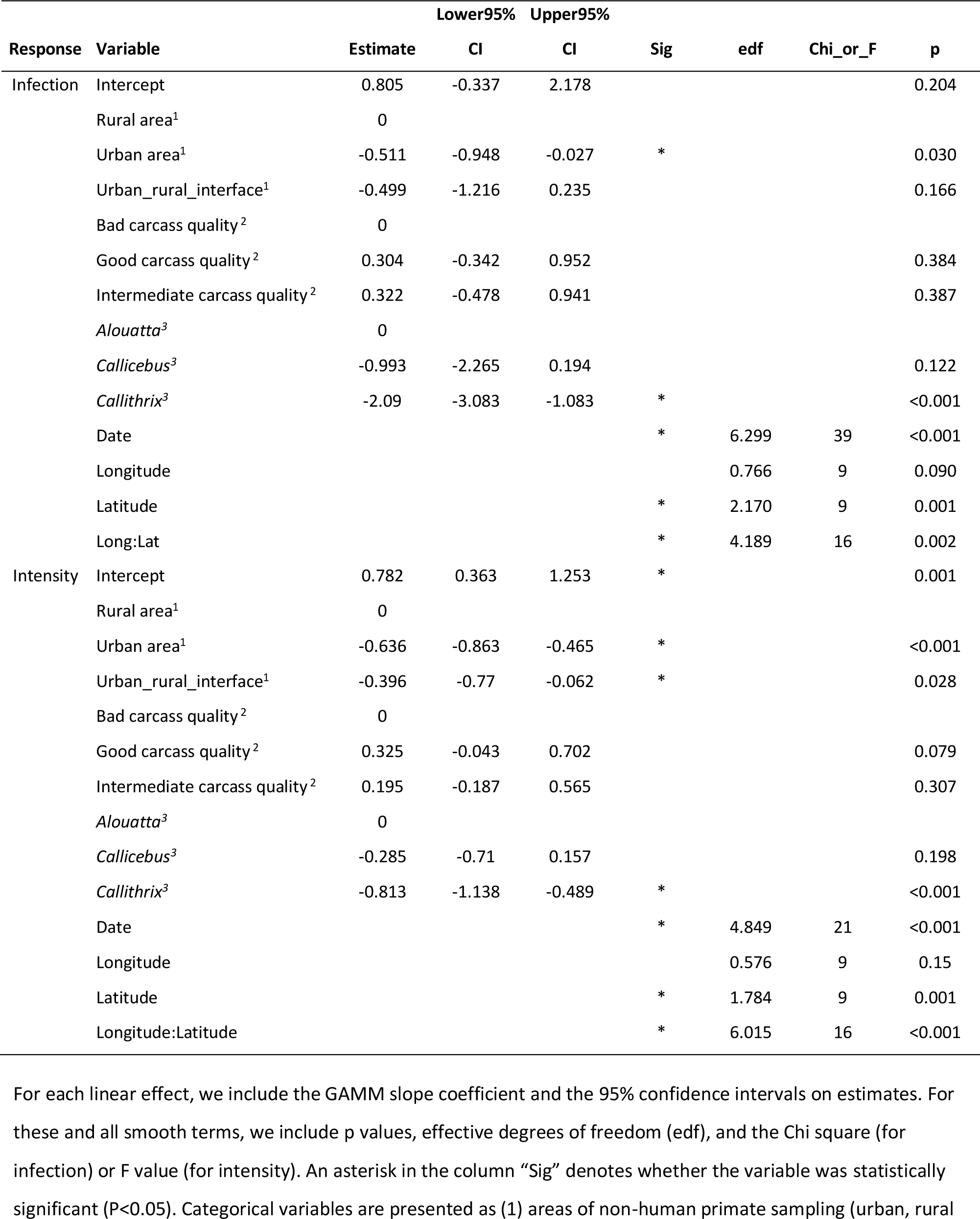
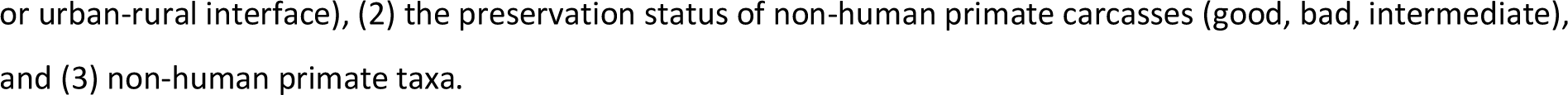
Effect estimates and P values for each of the terms in our generalized additive mixed models

**Table 3:** R values (correlations between predicted and observed values as a proportion of the model’s overall fit) for each of the fitted terms in our GAMMs.

| Response | Variable | R |
| --- | --- | --- |
| **Infection** | Area<sup>1</sup> | 0.0116 |
|  | NHP carcass quality<sup>2</sup> | 0.0015 |
|  | Taxon<sup>3</sup> | 0.0514 |
|  | Date | 0.5445 |
|  | Spatial effect | 0.0398 |
| **Intensity** | Area<sup>1</sup> | 0.1863 |
|  | NHP carcass quality<sup>2</sup> | 0.0175 |
|  | Taxon<sup>3</sup> | 0.1511 |
|  | Date | 0.2178 |
|  | Spatial effect | 0.1037 |

NHP: non-human primate. Variables are presented as (1) areas of non-human primate sampling, (2) the preservation status of non-human primate carcasses, and (3) non-human primate taxa.

First, we tested the hypothesis that the temporal pattern of the YFV epizootic in NHPs would mirror that of human cases, with peaks in NHPs preceding those of humans. However, the temporal patterns of total epizootics (Fig 2a) and human cases (Fig 2d) (based on data from the Ministry of Health) and the YFV infection in NHP studied here were roughly equivalent (Fig 2b), and the temporal smooth was both highly significant (P<0.001; Table 2) and accounted for a large proportion of the model deviance (R=0.54 and R=0.22; Table 3). We observed a strong temporal pattern in which YFV infection and intensity were higher during warm/rainy periods (Fig 2, Table 2), consistent with well-known seasonal patterns of YF (5). Broadly, both infection and mean intensity in our NHP samples were high at the start of 2017, and then dropped before increasing rapidly near the start of 2018. In NHP’s, both YFV infection and intensity peaked in early 2018, approximately the same time the human cases peaked, and then decreased extremely rapidly (Fig 2b-c). The penultimate positive NHP carcass was identified on 18^th^ March 2018, followed by a final positive carcass on April 13^th^ 2018 (Fig 2b). As noted above, one positive sample was identified on June 18^th^, 2020, but excluded from GAMM analyses.

**Figure 2:**
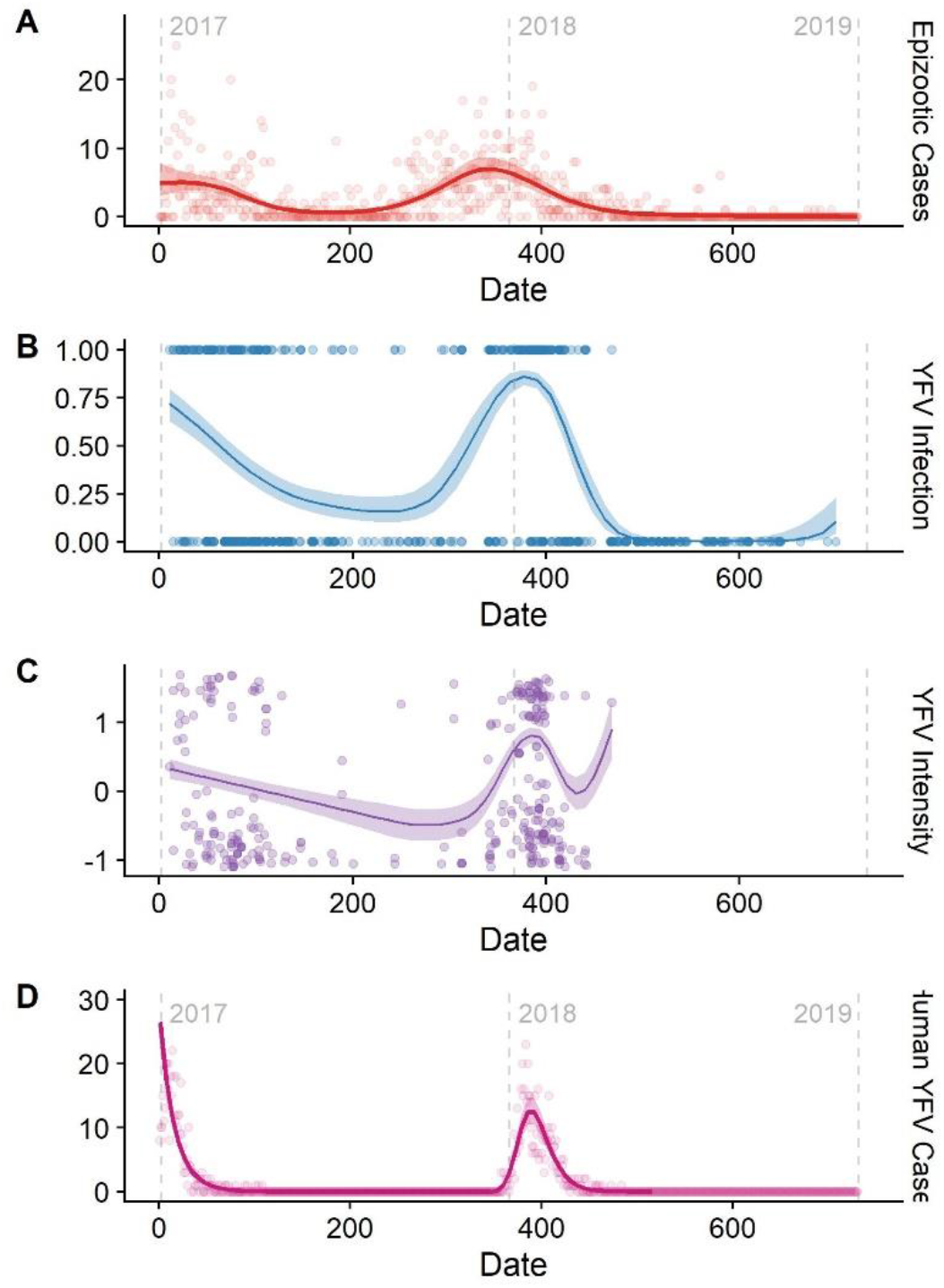
The temporal progression of (from top to bottom): (A) total epizootic cases, taken from the Brazilian Ministry of Health’s official yellow fever surveillance program; (B) predicted YFV infection in sampled primates (points indicating positive and negative samples); (C) intensity of YFV in infected primate samples and labeled in Panel B; and (D) total of human cases taken from Brazilian Ministry of Health’s official yellow fever surveillance program. Date is presented in “days after the start of 2017,” with the start of each year signposted by grey dotted lines. The fitted line is the mean fit taken from our GAMMs, using an adaptive smooth; the shaded area represents the 95% confidence intervals. YFV intensity was scaled to have a mean of 0 and a standard deviation of 1. Note: the upturn to the right of the intensity curve (Panel C) is likely an artifact of the single high-intensity point that was the final collected sample, and should be disregarded.

Second, we tested the hypothesis that the epizootic would be spatiotemporally structured and associated with particular land cover classes. There were strong, highly significant spatial patterns of YFV infection and intensity (p<0.001; Table 2) that accounted for substantial deviance (R=0.04 and R=0.1; Table 3). However, patterns for infection and intensity were less correlated in space than they were with time (Fig 3) and our GAMM model fits were not improved when we allowed the spatial patterns to vary between warm/rainy periods (ΔDIC<2), implying that spatial patterns were relatively similar in these periods. Spatial patterns of epizootics during the first (2017) and second wave of YFV (2018) were similar, implying that the drivers of these epidemics did not differ notably across epidemics. Although the intensity patterns were patchier and therefore more difficult to generalize, there was a band of low prevalence stretching across the central-west part of the state, with a relatively higher infection closer to the eastern part of Minas Gerais, where most human and NHPs cases had reported during outbreaks prior to the 2017-2018 outbreak (11,23,30,44). However, in this latest epidemic, human YFV cases were more concentrated in the northeast (2017) or in the central/southeast parts of Minas Gerais (2018), thus diverging somewhat for the foci of NHP infection (23,30).

**Figure 3:**
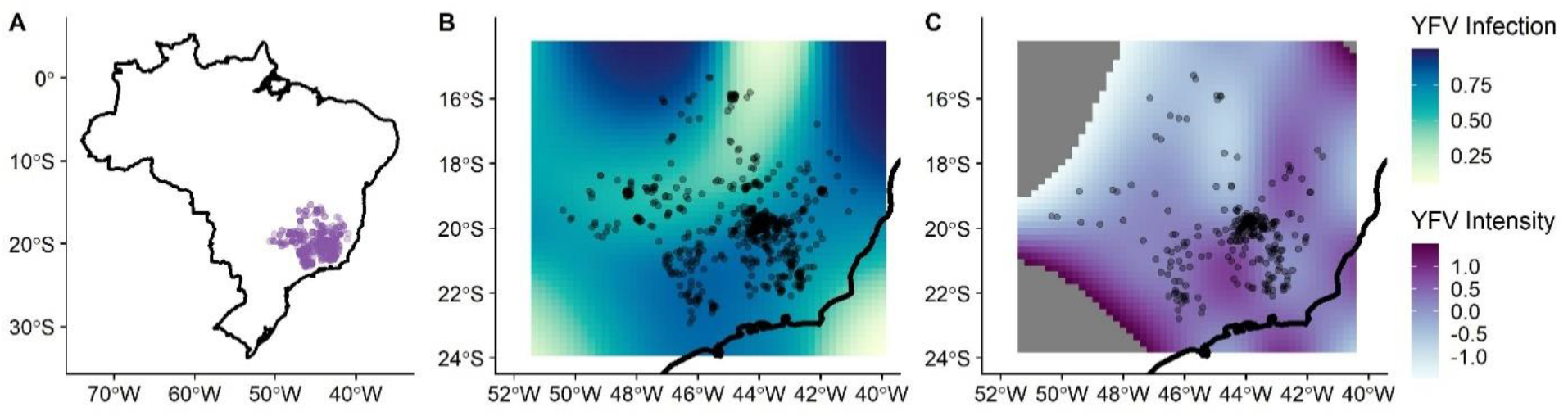
The geographic distribution of non-human primate carcasses (A); predicted YFV infection (B); and predicted mean YFV intensity (C). Panels B and C were taken from our GAMMs; darker colors represent greater predicted infection and intensity. The dark black line represents the coastline of Brazil. In panels B and C peripheral areas with very low or very high predicted intensity have been removed for plotting clarity (dark grey areas), due to a tendency for extreme estimates when extrapolating outside the limits of the data.

With respect to land cover, our GAMMs revealed that monkeys collected from the urban-rural interface had lower intensity (p=0.028), but not lower odds of infection (p=0.17), than those taken from rural areas (Fig 4; Table 2); samples from fully urban areas had both lower intensity (P<0.001) and probability of infection (p=0.03) than those taken from rural areas. The finding of higher YFV prevalence in rural areas is consistent with the sylvatic transmission cycle of the virus and likely reflects the distribution of key YFV vectors in the genera *Haemagogus* and *Sabethes* (3,11).

**Figure 4:**
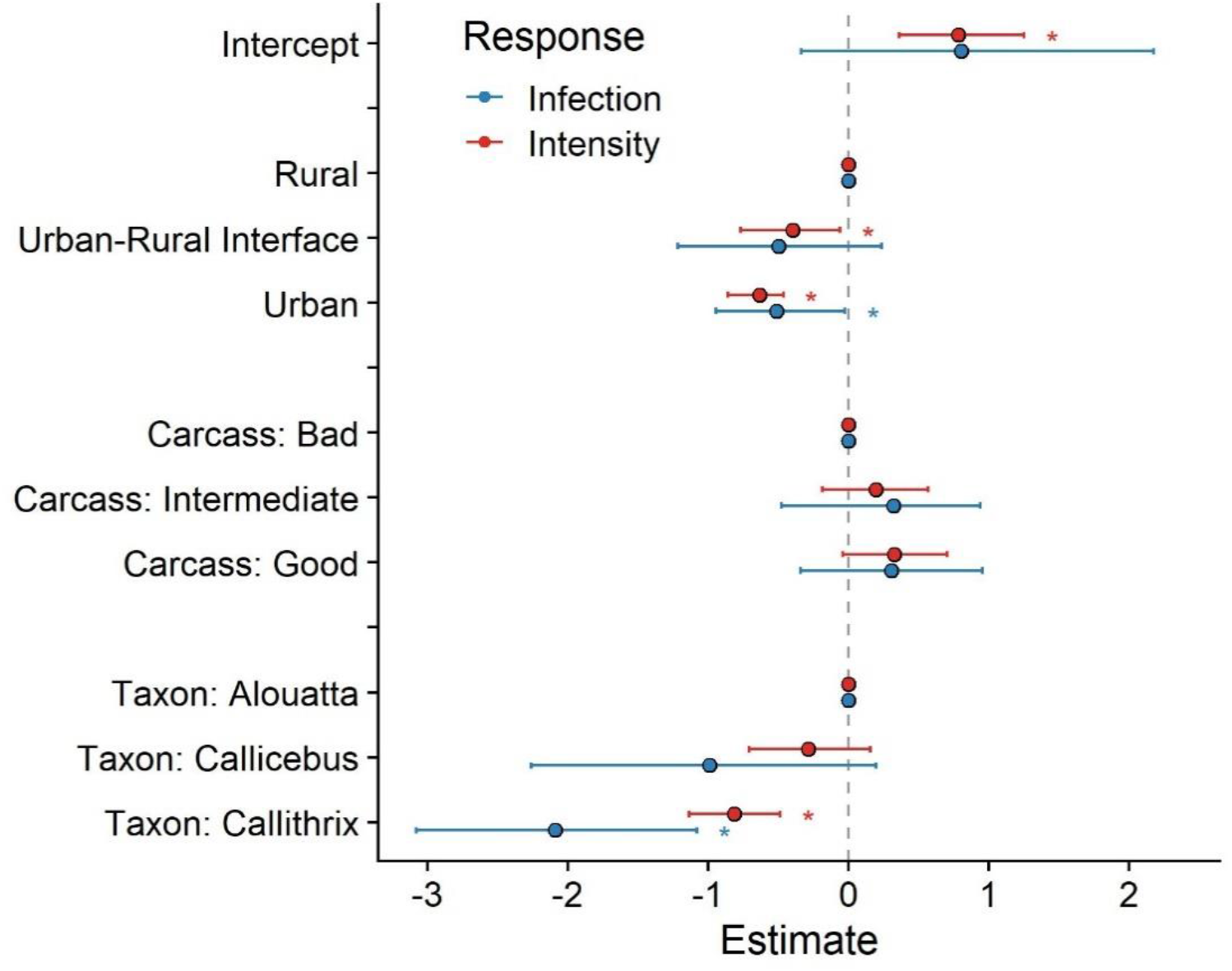
Fixed effect estimates taken from our GAMMs, for YFV infection (red) and intensity (blue), accounting for the nonlinear spatiotemporal variation seen in Fig 2-3. Dots represent the mean estimated effects; error bars are the 95% credibility intervals. Significant effects (p<0.05) are marked with an asterisk (*). “Rural”, “Carcass: Bad”, and “Taxon: Alouatta” were all the base levels for their factor; we display them as zero to allow comparison between these factor levels and the other factor levels in the effect.

Lastly, our GAMMs demonstrated that *Callithrix* had lower odds of infection and lower intensity than samples from the genera *Alouatta* and *Callicebus* (p<0.01). This pattern could be due to differential susceptibility of different to YFV infection (24–26), or to the higher percentage of *Callithrix* collected from urban areas, given that infection was shown to vary by land cover (3,11).

### Concordance of YFV Infection Prevalence and Intensity

We found a strong, positive association between the likelihood of infection (prevalence) and infection intensity (inverse Cq value representing concentration of viral genomes) in NHPs across time and space, with peak infection and intensity occurring in the warm/rainy season in eastern Brazil. While such a relationship might be expected in an ectothermic mosquito vector, where susceptibility and virus replication are both coupled to temperature (discussed below), it is not expected *a priori* in endothermic hosts. We hypothesize that this relationship may be driven by differences in the viral dose delivered to monkeys in the warm rainy season, either because NHPs receive more infectious bites during this season, or because individual mosquitoes deliver more virus per bite during this season, or both. The rationale for this hypothesis comes from the documented effects of varying virus dose in experimental arbovirus infections. Althouse and colleagues (50,51) as well as Fox and Penna have reported that experimentally increasing the dose of arbovirus delivered to an NHP or other host species leads to a higher peak viremia. Similarly, Hanley and colleagues (53) demonstrated that when mice were infected with different doses of Zika virus via different numbers of infectious mosquito bites, increasing dose resulted in higher peak viremia. Intriguingly, Hill and colleagues (54) found that during the 2016-2018 YFV outbreak in Brazil, the rate of YFV evolution is higher was faster the warm/season than the cool/dry season, a difference that they attributed to differences in virus infection dynamics within mosquitoes but that could also stem from variation in infection dynamics in NHP hosts.

It is known that the warm/rainy season favors the reproduction of the sylvatic vectors of YFV (55, 56) which in turn would result in higher vector-biting and therefore greater potential rates of NHP exposure to YFV Moreover, Chouin-Carneiro and colleagues (57) have shown that Zika virus infection and dissemination efficiency in Brazilian *Ae. aegypti* and *Ae. albopictus* is significantly greater in temperatures mimicking the Brazilian warm/wet season than temperatures mimicking the Brazilian cold/dry season. While the effects of temperature and humidity on vectorial capacity are quite complex (58), viral replication generally rises and extrinsic incubation period shrinks with increasing temperature up to some thermal limit. However, the conclusions above about the effects of temperature on virus infection, replication and dissemination are drawn from studies of *Aedes* mosquitoes, whereas the YFV outbreak under consideration here was sustained by *Haemagogus* (59) and possibly *Sabethes* mosquitoes. Unfortunately, the lack of colonized *Haemagogus* and the paucity of colonized *Sabethes* make it impossible to evaluate the impacts of temperature and humidity on the within-host dynamics of YFV replication in these genera. In lieu of such experimental studies, longitudinal surveillance of these sylvatic mosquitoes to quantify virus carriage and titer across seasons is needed. Moreover, our discovery of sustained, enzootic YFV transmission in Brazil likewise elevates the urgency of monitoring infection of sylvatic mosquitoes in this region.

## Acknowledgements

We do thank the Centers for Research in Emerging Infectious Diseases (CREID) (NIH 1U01AI15378), CREID Pilot Research Program, the Coordinating Research on Emerging Arboviral Threats Encompassing the Neotropics (CREATE-NEO), Laboratório de Zoonoses and Centro de Controle de Zoonoses da Prefeitura de Belo Horizonte, Secretaria de Estado de Saúde de Minas Gerais, colleagues from Laboratório de Vírus/UFMG, and Pró-Reitorias de Graduação, de Pós-graduação e de Pesquisa/Universidade Federal de Minas Gerais/Brazil. This study was developed with the participation of students from the Graduation Program in Microbiology of Universidade Federal de Minas Gerais, which is supported by Coordenação de Aperfeiçoamento de Pessoal de Nível Superior - Brasil (CAPES - 88882.348380/2010-1).

## Funding

We thank the Centers for Research in Emerging Infectious Diseases (CREID) and CREID Pilot Research Program 2021 (grants 1U01AI15378, and X.0217530, respectively, by the National Institutes of Health (NIH). KAH and NV research is supported by Centers for Research in Emerging Infectious Diseases [CREID], “The Coordinating Research on Emerging Arboviral Threats Encompassing the Neotropics [CREATE-NEO]” grant U01 AI151807 by NIH. GFA, DB, and CJC were supported by NSF BII 2021909 and NSF BII 2213854. BPD is a CNPq/MCTI research fellow. This study was developed with the participation of students from the Graduation Program in Microbiology of Universidade Federal de Minas Gerais, which is supported by Coordenação de Aperfeiçoamento de Pessoal de Nível Superior - Brasil (CAPES) (grant 88882.348380/2010-1). Scholarships for students were provided by FAPEMIG, CAPES (001), PRPq-UFMG/PIBIC-CNPq. The funders had no role in the design of the study, collection, analyses, or interpretation of data, writing of the manuscript, or in the decision to publish the results.

## Notes

https://github.com/viralemergence/minas-gerais-yfv

### Competing Interest Statement

The authors have declared no competing interest.

